# Botanicals impact the bifidogenic effect and metabolic outputs of *in vitro* fiber fermentation by gut-derived microbiota in individual-specific ways

**DOI:** 10.1101/2023.04.17.537114

**Authors:** Dane G. Deemer, Noah Voreades, Peter A. Bron, Stephen R. Lindemann

## Abstract

Fortification of products frequently consumed by a large proportion of society provides an attractive strategy to close the “fiber gap” and may have the potential to concomitantly reverse the detrimental health effects exacerbated by our modern diets. Besides prebiotic fibers, products can contain other functional components, e.g. botanicals. However, most studies have investigated functional components in isolation. The impact of other components present in functional product blends on the bifidogenic effect typically exerted by prebiotic fibers are largely unexplored. Here, we investigated the fiber and botanical blends included in OLIPOP, a functional soda, in an *in vitro* gut fermentation model. Our data revealed that the blend of inulins and resistant dextrins promoted growth of bifidobacteria across gut microbiota from four donors, even those with small initial populations. In addition, botanicals interacted with fiber fermentation in donor-specific ways, in some cases strongly enhancing fermentation rate and production of short-chain fatty acids.

**Graphical Abstract:** 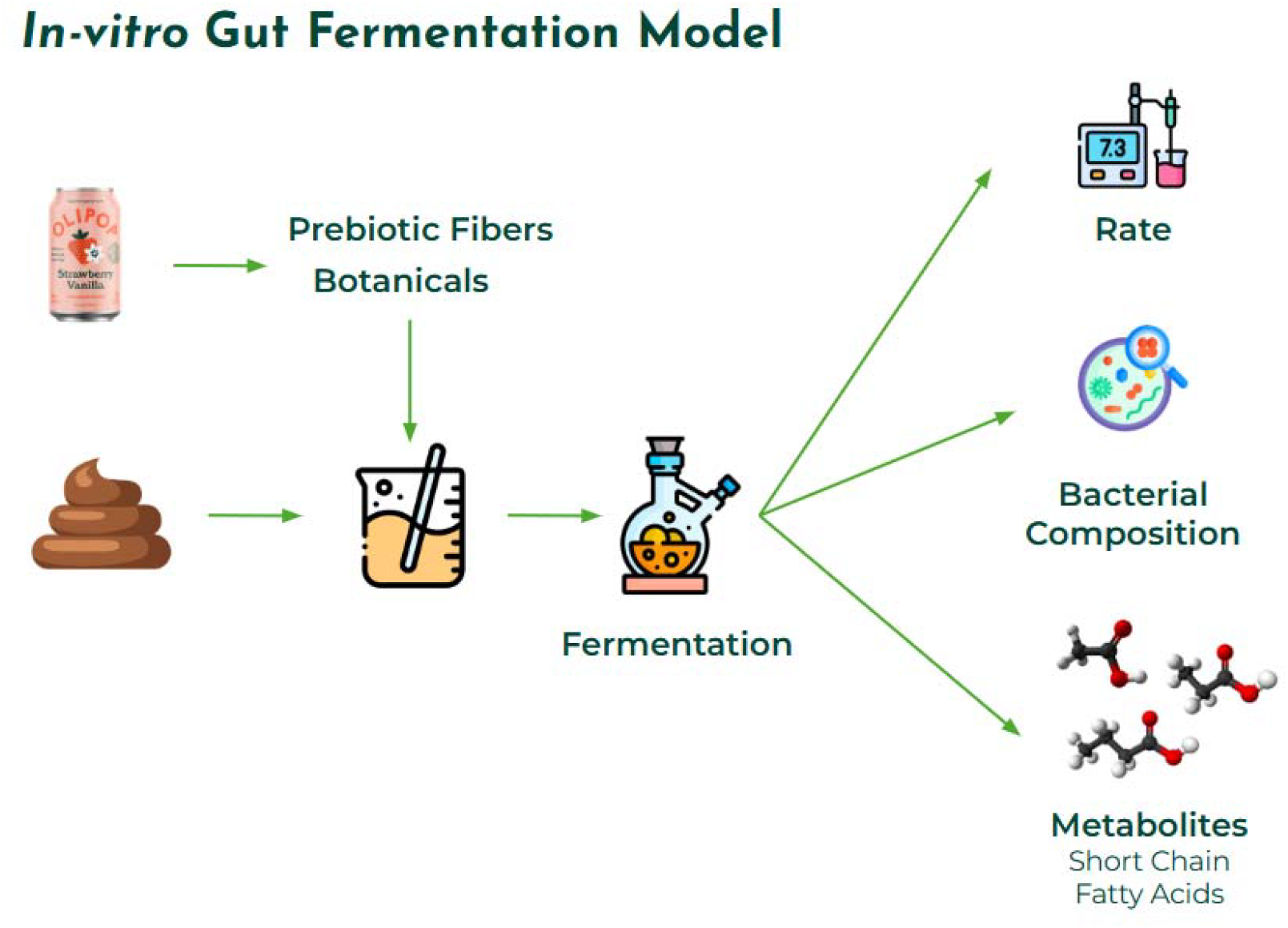

## 1 Introduction

Carbohydrate intake in modern Westernized diets has significantly shifted in fiber content and composition compared with ancestral human diets, both with hunter-gather and agricultural traditions (De Filippo *et al*., 2010; Schnorr *et al*., 2014). Current hunter-gatherer populations, such as the Hadza of Tanzania, typically consume upwards of 100g of fiber per day and approximately 70% of their caloric intake is plant-derived (Smits *et al*., 2017). By contrast, the modern-day population overwhelmingly consumes easily-digested carbohydrate types, such as simple starches (54%) and sugars (36%). Currently, Westernized diets are associated with high intakes of saturated fat and sucrose and are low in total fiber intake. Collectively, the consumption of a Western type diet represents a growing health risk for metabolic diseases such as diabetes and diseases of the gut, such as inflammatory bowel disease (Statovci *et al*., 2017). Other literature describes that Western diets and lifestyles are associated with cancer and less diverse gut microbiomes (O’Keefe *et al*., 2015; Vangay *et al*., 2018). Conversely, mounting scientific evidence suggests that increasing total fiber intake is beneficial both for supporting digestive, gut microbiome and overall health (Makki *et al*., 2018; Reynolds *et al*., 2019). Specifically, in the U.S., the Academy of Nutrition and Dietetics recommends that higher daily dietary fiber intakes may reduce risk of multiple chronic diseases, including type 2 diabetes, cardiovascular disease, and some cancers, and lower body weight (Quagliani and Felt-Gunderson, 2016). Consequently, the Academy suggests a minimum daily fiber intake of 14 g per 1,000 kilocalories for adult women and men, equaling 25 and 38 g/day, respectively (Dahl and Stewart, 2015).

Despite changes to nutrition policy, governmental reimbursement of healthier foods, and insurance programs incentivizing healthy food purchasing, modern societies fail to consume sufficient total fiber and diversity of fiber types on a daily basis. Furthermore, modern consumers are choosing increasingly restrictive dietary patterns (gluten-free, grain-free, wheat-free, etc) which may be leading to reduced daily fiber intake (Quagliani and Felt-Gunderson, 2016). Unfortunately, daily nutrition intake surveys analyzed from National Health and Nutrition Examination Survey (NHANES) cycles 2013 – 2018 indicate Americans on average only consume between 9g-10g/1,000 kcal of total fiber per day, which translates to less than 1 in 10 of the population meeting minimum suggested fiber intake levels. This shortfall between recommended and actual fiber intake is commonly referred to as the “fiber gap.” Fiber fortification of foods and beverages can potentially bridge this gap and may, in this way, contribute to reduced incidence of chronic disease in industrialized urban societies. Even the CODEX Alimentarius Commission has aligned its definition to support bridging the fiber gap (Jones, 2014). To this end, one rapidly growing category in the beverage sector is functional soda, which aims to provide consumers with healthier alternatives to traditional soda, often by reducing sugar and adding various prebiotic fibers. Such approaches hold promise for modulating population-scale fiber intake because the selection of a high-fiber alternative with a similar sensory profile within the same product category often requires relatively minimal behavioral adaptation. This ease of adoption by consumers may explain the rapid growth of this product category. Moreover, success in this product category likely serves as a model for fiber augmentation/sugar replacement strategies in other categories.

The mechanisms by which fibers exert beneficial health-promoting properties are multi-factorial, but include increased production of short-chain fatty acids (SCFAs), stimulation of beneficial microbial taxa (e.g. bifidobacteria), reduction in the production of nitrogenous and otherwise pro-inflammatory metabolites, ammonia, amines, and some phenolic compounds (Makki *et al*., 2018). SCFAs produced by gut bacteria through fiber fermentation support healthy gut epithelial barrier and immune function. Beyond the role of SCFAs in gut barrier integrity, dietary fibers interacting in the digestive tract mechanically stimulates the gut epithelium to secrete mucus. Depletion of dietary fiber consumption compromises gut barrier integrity, which increases the risk for acute infection and chronic disease (Desai *et al*., 2016). Moreover, high fermentable fiber consumption provides substrates for the growth and maintenance of beneficial microbial populations in the colon. However, leveraging fiber supplementation to bridge the fiber gap requires a more thorough mechanistic understanding of the relationship between microbiome compositional changes, metabolite production, and human physiology.

Dietary fiber is defined by the FDA in 21 C.F.R.§101.9(c)(6)(i) as “non-digestible soluble and insoluble carbohydrates (with 3 or more monomeric units), and lignin that are intrinsic and intact in plants; or isolated or synthetic non-digestible carbohydrates (with 3 or more monomeric units) determined by FDA to have physiological effects that are beneficial to human health.” Some dietary fibers are termed “prebiotic” according to the International Scientific Association for Probiotics and Prebiotics (“ISAPP”) definition (Gibson *et al*., 2017), which means that they are selectively utilized by gut microbes; the concept of a prebiotic has recently been expanded to include diverse other molecule types (e.g. polyphenols). Fibers that are also designated prebiotics are often termed “prebiotic fibers” on consumer products. Thus, “dietary fiber” is a heterogeneous category of plant carbohydrates, having wildly diverse physical and chemical structures and associated physiochemical properties (solubility, viscosity). It is very likely that specific fiber structures exert divergent impacts directly on the host and indirectly via its gut microbiota. For example, even though they are all glucans, a set of commercial mixed-linkage □-glucans, resistant dextrins, and polydextroses produced as supplemental fibers for food products fermented to distinct metabolic and microbial outcomes; however, some taxa exhibited specificity for some glucans across donors (Romero Marcia *et al*., 2021). Successful utilization of fibers and prebiotics in designing fiber-supplemented food and beverage products to improve health via the microbiome at population scales requires much greater insight into such mechanistic linkages among fine fiber structure and microbiota.

Importantly, fiber fermentation *in vivo* does not occur in a vacuum; fibers are fermented in the context of many other compounds that might impact their utilization by microbes and metabolic fate in the colon. Some functional foods and beverages deliberately employ plant extracts (hereafter, “botanicals”) and probiotics, prebiotics or postbiotics, with the goal of enhancing their health benefits. Increasing our understanding of a botanical extract’s influence on the microbiome and human health is important given their inclusion in numerous dietary supplements or functional food and beverage products in recent years (https://tinyurl.com/krmet4vs). Furthermore, botanical extracts being included in dietary supplements or functional food and beverage products is not unique; frequently, companies are creating multi-ingredient products including botanicals and prebiotic fibers. Although mechanistic and human health outcomes are widely studied for prebiotic fibers, there is a paucity of scientific literature assessing the effects of botanical extracts on the gut microbiome or broader human health. Moreover, to the best of our knowledge, no microbiome studies have been performed elucidating the combined interaction effects of botanical extracts and prebiotic fiber within a final product formulation. Developing mechanistic and human clinical trial datasets on a product’s functional blend to complement research on isolated ingredients is necessary to understand microbiome and beneficial human health effects.

*In vitro* batch culturing systems mimicking fermentation by colonic microbiota are a valuable tool to screen the impact of product formulations on the microbiome prior to the execution of larger clinical studies (Yao, Chen and Lindemann, 2020). Here, we assessed the impact of the prebiotic fibers and botanicals in OLIPOP, a functional soda, on gut microbiome composition and functionality, by employing an anaerobic batch culturing system inoculated with fecal slurry from 4 donors. Through the inclusion of study arms containing OLIPOP formulations with and without added botanical extracts, the interaction impacts of these important dietary components could also be investigated.

## 2 Materials and Methods

### 2.1 Donor Selection

Fecal donors were recruited for this study under oversight from the Purdue University Institutional Review Board under protocol IRB-2020-1650. Four donors were selected that satisfied the following criteria: 1. between 18-39 years old, 2. having a normal or overweight BMI (18.5 < BMI < 30), 3. having been consuming their normal diet for two weeks, 4. have not taken any antibiotics within the last 12 months, 5. have not taken fiber, prebiotic, or probiotic products within the last 3 months, and 6. are not heavy alcohol drinkers (consuming 5 or less alcoholic drinks per day). Donors selected had no history of gastrointestinal or chronic metabolic diseases and had not had a major gastrointestinal surgery in the past 5 years or a major bowel resection ever. Fecal samples were collected from two male and two female donors aged 20-39. Both males (Donors 1, 38 years old and 2, 27 years old) and one female (Donor 4, 28 years old) reported diets consistent with omnivory, whereas the other female (Donor 3, 32 years old) reported a vegetarian diet.

### 2.2 Fermentation Substrates

Carbon sources were provided by OLIPOP, Inc. shipped on ice, and immediately placed at 4°C. Two different carbon sources were used to supplement the medium used during the fecal fermentations: a blend of fibers (cassava root fiber, chicory root inulin, and Jerusalem artichoke inulin; “the OLISMART fibers”) reconstituted in sterile water, and the same reconstituted OLISMART fibers combined with a mixture of botanicals (extracts from nopal cactus, marshmallow root, calendula flower, and kudzu root). We maintained the concentrations of fibers and botanicals equivalent to those found in any complete OLIPOP soda product found at retail - 9 g of dietary fiber per one 355 ml can. This equated to a total substrate loading of 0.025g/ml, or 2.5% carbohydrate (w/v).

### 2.3 *In vitro* Fermentation

One liter of the basal fermentation medium contained the following substances: 0.001g resazurin, 0.10g Na_2_SO_4_, 0.40g urea, 0.45g KCL, 0.468g NaH_2_PO_4_, 0.47g NaCl, and 0.865g Na_2_HPO_4_ and autoclaved (121°C for 30 minutes). Heat sensitive compounds CaCl2, MgCl_2_, 1mL cysteine hydrochloride (0.25g/L) and 1mL of 1000X P1 metal solution were added via 0.22μM filter sterilization. Per liter, the P1 metal solution was composed of 34.26 g H_3_BO_3_, 4.32 g MnCl_2_ · 4H_2_O, 0.315g ZnCl2, 44mg Na_2_MoO_4_ · 2H_2_O, 3mg CuSO_4_ · 5H_2_O, 12.15 mg CoCl_2_ · 6H_2_O, 259mg NiCl_2_, 0.28ml EDTA (10mM),1ml FeCl_3_ · 6H_2_O (3.89 mg/ml in 0.1MHCl). The media was fortified with 200μM sterilized amino acids (10μM each) and 1% (v/v) ATCC vitamin supplement (ATCC MD-VS; Hampton, NH) and finally pH controlled at 7.0. The amino acid mixtures (200μM final concentration in media) contained 20 proteinogenic amino acids, including: alanine, glycine, isoleucine, leucine, proline, valine, phenylalanine, tryptophan, tyrosine, aspartic acid, glutamic acid, arginine, histidine, lysine, serine, threonine, cysteine, methionine, asparagine and glutamine at final concentrations of 10μM each.

The base media was used as a blank, with either OLISMART (prebiotic fibers and botanicals mentioned above) or OLISMART excluding botanicals solutions added for experimental conditions. All fermentations were performed in an anoxic chamber (Coy Laboratory Products Inc., Great Lake, MI, USA) supplied with nitrogen, carbon dioxide, and hydrogen (90%, 5%, and 5% respectively). The media were placed in the anoxic chamber 24 hours before the fermentation in order to reduce oxygen levels.

Fecal samples were obtained from donors in the early morning using a custom fecal collection kit. Fecal samples were collected in sterile 50mL Falcon tubes and stored immediately on ice. Fecal inocula were anoxically prepared as previously described (Yao, Chen and Lindemann, 2020) with the following modifications. Briefly, fecal aliquots were diluted 1:5 in medium and homogenizing via rapid pipetting and vortexing. The fecal slurry was further diluted in a 1:20 ratio with the corresponding medium type (no additions, fibers, fibers and botanicals) and poured over 4 layers of cheesecloth to remove large particles, yielding a final fecal slurry consisting of a 1:100 dilution of fecal material. 5.0 mL of the prepared fecal slurry was placed in 15 mL Balch tubes and sealed with a butyl rubber stopper plus an aluminum seal. The Balch tubes were placed in an incubator at 37°C, shaking at 150 rpm. Fermentation cultures were harvested in triplicate for each of the 3 media at timepoints 2, 4, 8, 12, 18, and 24 hour. Two aliquots of 1.5mL were used from each tube for downstream SCFA analysis and DNA extraction, and pH was measured on the remaining spent media.

### 2.4 DNA Extraction and Sequencing

Upon donation of fecal samples on ice, three tubes were filled with raw fecal material (∼500 mg), and three more tubes were filled with 1.5mL of 1:100 feces to medium mixture and immediately placed at -80 ^o^C. During the fermentation, three fermentation tubes were sacrificed for each condition at each timepoint, and the product was immediately placed at - 80°C in 1.5mL aliquots. Raw feces, baseline blanks (medium with fecal slurry at the initiation of the experiment), and fermentation products (fiber, fiber and botanicals) at each timepoint were then sent to Diversigen™ for DNA extraction via PowerSoil Pro (Qiagen), automated for high throughput processing on the QiaCube HT (Qiagen), using Powerbead Pro Plates (Qiagen) with 0.5mm and 0.1mm ceramic beads. Samples were quantified with Quant-iT PicoGreen dsDNA Assay (Invitrogen). Libraries were prepared with a procedure adapted from the Illumina DNA Prep kit. For shallow sequencing, libraries were sequenced on an Illumina NovaSeq using single-end 1x100 reads. DNA sequences were filtered for low quality (Q-Score < 30) and length (< 50), and adapter sequences were trimmed using cutadapt.

### 2.5 Metagenomic Analysis

A conda virtual environment was created on Purdue University’s Bell cluster (CentOS 7) with Python version 3.7.10. All jobs were run using Dell compute nodes with two 64-core AMD Epyc 7662 “Rome” processors and 256 GB of available memory. Sequences were downloaded as raw FASTQ files from Diversigen and FastQC (v0.11.9) was used to analyze initial metagenomic reads and determine if trimming and filtering were required. Reads were taxonomically assigned using MetaPhlAn (v3.0.7) and PhyloPhlAn (v3.0.2), which use reference-based algorithms to taxonomically identify reads up to the species level. To generate raw abundance calculations of taxonomic representation per sample, reads were summed at the highest specificity of classification. Relative abundances were then computed using total library size as a normalization factor for each sample. Reads without a taxonomic classification were removed from further analysis.

Alpha and Beta diversities were calculated by phyloseq (v1.30.0) using relative abundances of taxonomically classified reads as input. Beta diversity was calculated using the Bray-Curtis, Jaccard, and UniFrac methods. HUManN (v3.0) was used to profile the abundance of microbial pathways in the sequenced communities to determine metabolic potential.

### 2.6 Short-Chain Fatty Acid (SCFA) Analysis

SCFAs were measured at each of the timepoints in triplicate using a gas chromatograph (Nexis GC-2030, Shimadzu) with a flame-ionization detector and Stabilwax-DA column (Crossbond Carbowax polyethylene glycol; Restek, Inc.). The parameters of the analysis were as follows: 30m long by 0.25mm diameter column with 0.25μM film thickness, 230 °C injector temperature, and 100°C initial oven temperature with a max column temperature of 260 °C. 0.5μL of sample were injected and measured for 23.43 minutes in triplicate for each sample. External standards were used to back-calculate sample concentrations of acetate (Thermo Fisher catalogue number A38S), propionate (A258), butyrate (AC108111000), isobutyrate (AC122520250), and isovalerate (AAA18642AU).

Frozen aliquots consisting of 1.5mL of fermentation product were thawed at room temperature and spun at 13,000 ×*g* for 10 minutes. Supernatant was mixed in a 4:1 ratio with 4-methylvaleric acid (Thermo Fisher catalogue number AAA1540506) in 6% *v/v* phosphoric acid and copper sulfate pentahydrate; these additional products added to the sample made up the internal standard.

### 2.7 Statistical Analyses

An analysis of variance (ANOVA) was independently run on acetate, butyrate, and propionate concentrations with factors including donor, timepoint, and media source to test main effects. The base R (version 2021.09.1) function *aov()* was used to determine the main effects each independent variable had on acid production. 2-group t-tests through base R (function *t*.*test()*) were used to test for significant acid production differences at specific timepoints for each donor individually. Pairwise multiple testing and p-value adjustments were performed using Tukey’s honest significant difference (R stats library; function *TukeyHSD()*) to determine statistically significant acid production for each media type across each donor. Significant alpha diversity changes were also investigated with factors including Donor, Time, and Sample using *aov()* and *TukeyHSD()* R functions.

Metagenomic reads were quantified at the taxonomic level of family and subjected to ANOVA with read count as the dependent variable and media condition and donor as the interacting independent variables. Each donor’s abundance data was independently analyzed for condition effects on family-level read-count information. The base R function *aov()* was used to model the effects and *TukeyHSD()* (stats library) was used with a confidence level of 0.95 to determine significant differences across each group.

The R session, code, and input files required to replicate each of our plots, as well as raw input are available as supplementary information.

## 3 Results

### 3.1 OLISMART fibers are strongly fermented by gut microbiota but fermentation rate responses to botanicals are donor-specific

All *in vitro* fermentations containing OLISMART fibers proceeded rapidly, as measured by medium acidification, and reached terminal acidities (pH 3.5-4.0) within 12 hours of inoculation for donors 1 and 2, whereas pH continued to modestly decline in cultures from donors 3 and 4 for the duration of the experiment (Figure 1). Interestingly, the effect of botanicals on fermentation rate was donor-specific; botanicals substantially increased the fermentation rate in donor 1 and donor 3 cultures, had little or no impact on donor 2 culture rates, and significantly slowed acid generation in donor 4 cultures. In contrast, control fermentations to which no carbon source was added did not display a significant acidification, with pHs remaining between 6-7 for the entire duration of the experiment (data not shown). Thus, although fermentative responses to OLISMART fibers by gut microbiota were very similar across donor microbiota, the interactions of fibers with botanicals were determined by donor context.

**Figure 1:**
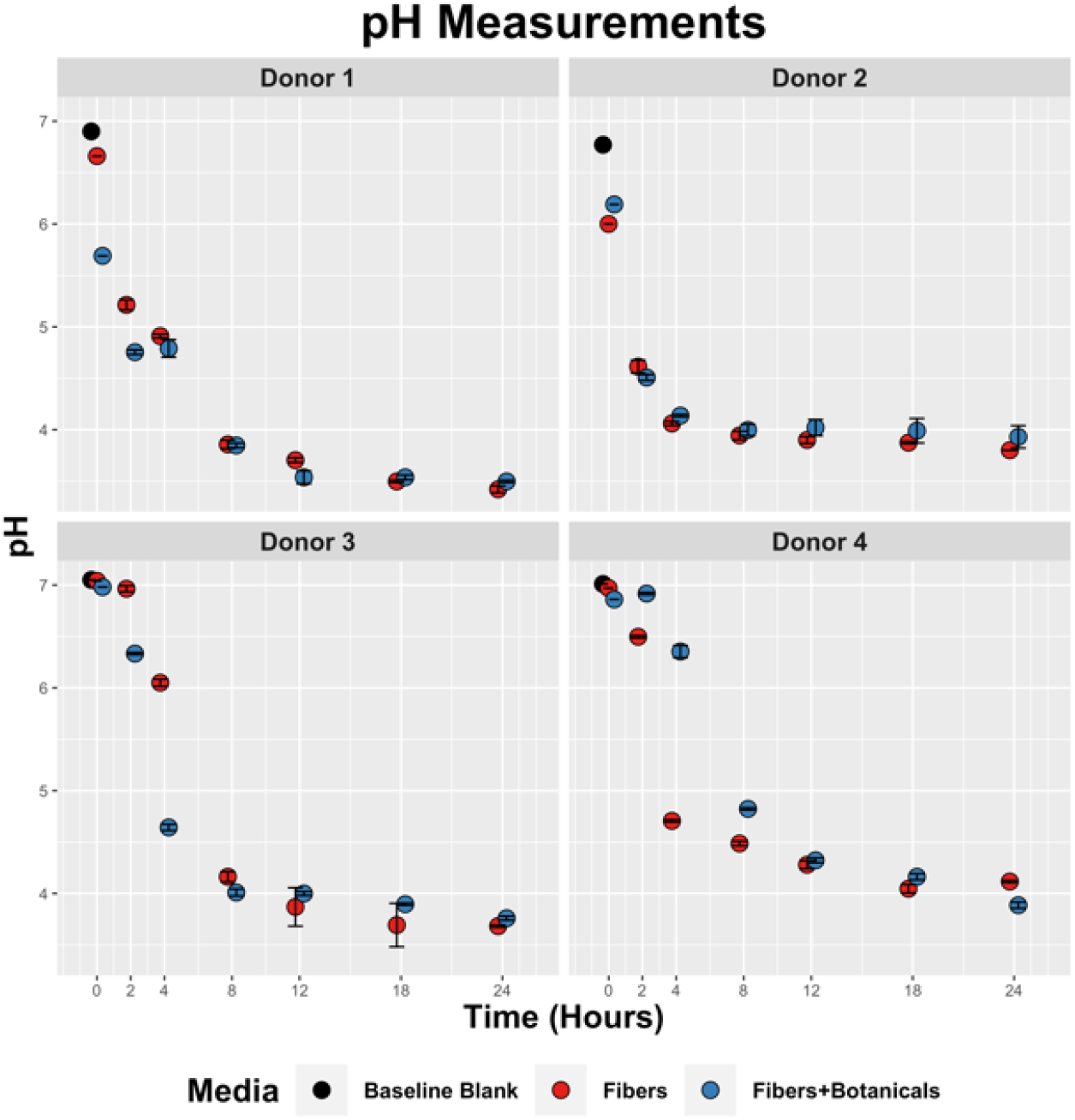
Temporal pH reduction by fecal slurries containing OLISMART fibers or OLISMART fibers supplemented with botanicals.

### 3.2 Short-chain fatty acid production levels are donor and botanical specific

Fermentation of OLISMART fibers across all donors was strongly acetogenic, approaching or exceeding 60 mM by the end of the fermentation (Figure 2A). The effect of botanicals on fiber fermentation was donor specific, generally in ways that reflected overall acid production trends; acetogenesis of donor 1 and donor 3 microbiota were significantly stimulated by botanicals at 12 hours (p < 0.05) while donor 4’s microbiota was not significantly affected. In contrast, acetogenesis by donor 4’s microbiota was strongly delayed by botanicals for the first 18 hours, thereafter showing significant botanical stimulation of acetate at 24 hours (p < 0.05); these communities produced no meaningful amounts of acetate until 12 hours post-inoculation. Interestingly, acids were still being produced in these communities (see below); the pH of these cultures dropped below 5 by the 8-hour time point. Further, the acetogenesis rate in botanical-amended cultures increased with time, until the end of the experiment. These data suggest substantial rewiring of the metabolism of OLISMART fibers by the donor 4 microbiota, and more minor influences on the donor 1 and 3 communities.

**Figure 2:**
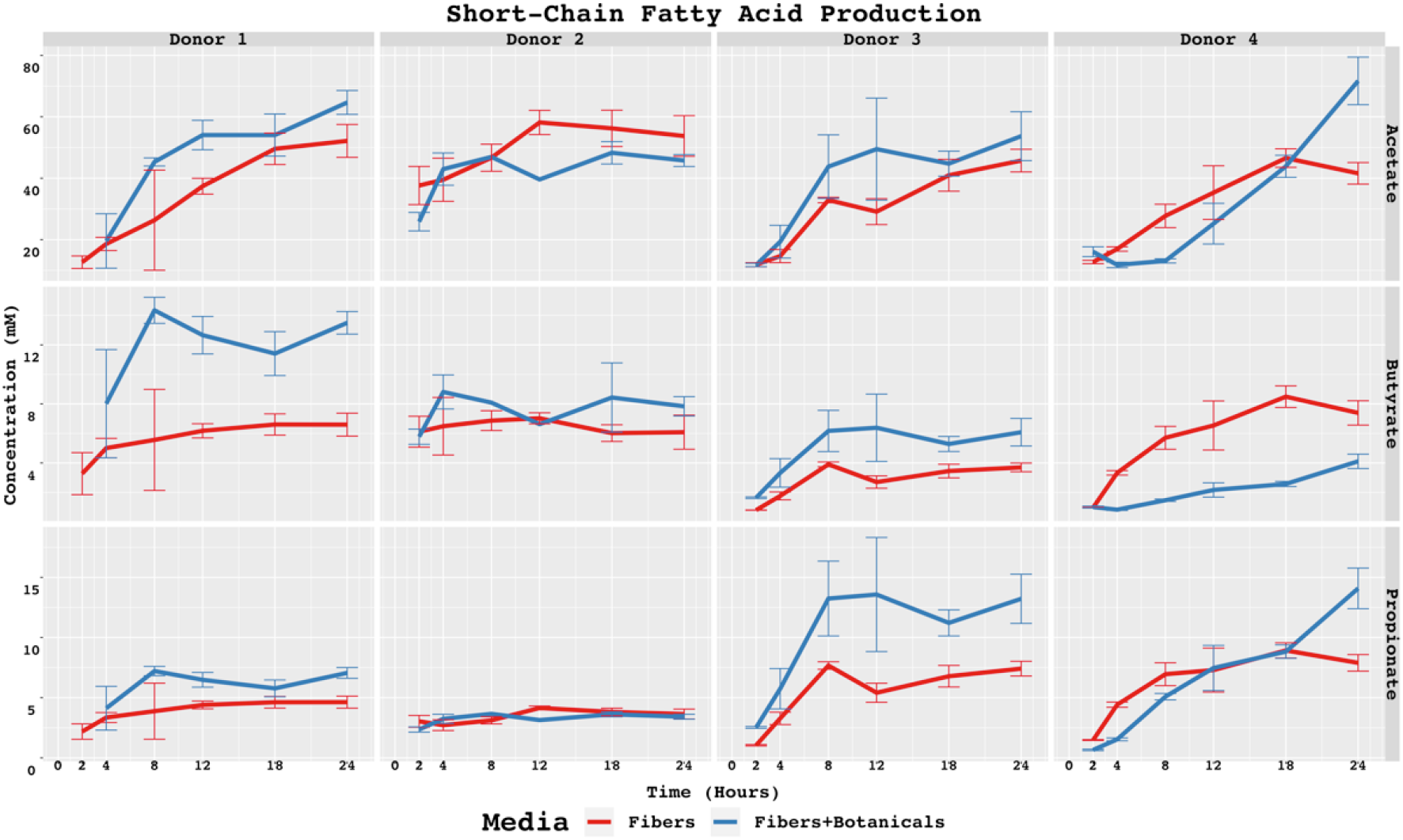
SCFAs concentrations in baseline controls and fiber fermentations with or without added botanicals. Statistically significant differences are calculated by pairwise t-tests with p < 0.05. Symbol style: p-value < 0.05 (*). Asterisk (*) above specific timepoints (x-axis) indicates statistical significance for timepoint; asterisk (*) in top right corner of panel indicates statistical significance at every timepoint for the donor-acid combination.

Unlike acetate production, production of propionate and butyrate from fermentation of OLISMART fibers was strongly donor- and botanical-dependent. Donor 1’s microbiota produced far more butyrate than propionate, and both levels appeared enhanced by the presence of botanicals (Figure 2; butyrate: ∼14 mM with and ∼7 mM without, p < 0.05; propionate: ∼7 mM with and ∼4.5 mM without, p < 0.05). Isobutyrate was also detectable only in donor 1 cultures with botanicals, which was also true for isovalerate (with the exception of t=2h, where this latter metabolite was also found in fermentations lacking botanicals). By contrast, fermentations containing donor 2 microbiota produced modestly more butyrate than propionate, and the production rates and amounts of these metabolites were similar with and without botanicals (p > 0.05 for all timepoints except propionate at 12 hours [likely due to limited number of replicates]); isovalerate and isobutyrate were undetectable in these cultures (data not shown). Donor 3’s microbiota were substantially more propiogenic than butyrogenic when OLIPOP fibers were fermented alone, and produced both significantly more butyrate (∼6 mM vs. ∼ 4 mM) and propionate (upwards in some cases of 15 mM vs. a maximum of ∼8 mM) when botanicals were present (Figure 2). ANOVA using acid concentration as the dependent variable revealed significant main effects of media, timepoint, and acid when comparing donor 3 butyrate and propionate SCFA data (p < 0.01 for all independent variables). Again, donor 4’s microbiota showed strong metabolic rewiring with botanicals; though slightly more propiogenic than butyrogenic without botanicals, when botanicals were added the production of butyrate dropped significantly (p < 0.05 for all timepoints except t = 2h) and propionate production was delayed, though it reached higher substantially concentrations at the end of the fermentation compared with fibers alone. We hypothesize that much of the SCFA production difference in donor 4 cultures was due to the accumulation of lactate, a common metabolic product of bifidobacteria, which was not measured in the study but is consistent with the magnitude of observed bifidobacterial blooms (similarly with acetate). Interestingly, donors that were strongly responsive to botanicals were the only ones in which BCFAs (isobutyrate, isovalerate) could be detected (data not shown), with maximum isobutyrate and isovalerate levels of 0.10μM and 0.30μM for Donor 1 and values of 0.11μM and 0.18μM for Donor 2, respectively.

#### 3.3 Fermentation of OLISMART fibers is bifidogenic across donors but the extent is individually modulated by botanicals

Although the microbial community composition responses to fermentation were individual and depended somewhat upon initial community composition (Figure 3), bifidobacteria expanded dramatically in relative abundance across all four donor populations during fermentation of OLISMART fibers either with or without added botanicals (p < 0.001, ANOVA using family read count as dependent variable and media type as independent variable). Metagenomic read counts generated for the *in vitro* fermentations revealed that across donors, members of phylum Actinobacteria, or more specifically of the family Bifidobacteriaceae significantly (p < 0.001, ANOVA) bloomed when grown on OLISMART fibers, with and without botanicals (Figure 4A). This rapid relative growth of bifidobacteria came at the expense of diverse members of Bacteroideaceae, Lachnospiraceae, and Ruminococcaceae in all donors, as well as Akkermansiaceae specifically for donor 4 (Figure 4B) (p < 0.001, ANOVA). It should be noted that reduction in the relative abundances of microorganisms does not imply that their absolute numbers are reduced, but rather that they are not growing sufficiently rapidly to keep up with the average growth rate of the community. Thus, these data should be interpreted that the growth rate of the bifidobacteria was substantially greater than other taxa in the community, regardless of the starting bifidobacterial population size. Not surprisingly, such strong expansion in one taxonomic group inherently led to significant (p < 0.001, ANOVA) time-dependent reduction in alpha diversity (a combination of species richness and evenness) over the 24-hour fermentation, as small populations of organisms fall below the limit of detection (reducing observed richness of the community given identical sampling effort) and the evenness of the community is reduced by the bloom in a single taxon (Figure 5). Notably, this reduction in alpha diversity was seen across all donors and no significant donor effects were observed (p > 0.1, ANOVA + Tukey HSD test). These data suggest that OLISMART fibers are strongly selective for bifidobacteria *in vitro* and this effect is observed even when initial endogenous populations of bifidobacteria are small.

**Figure 3:**
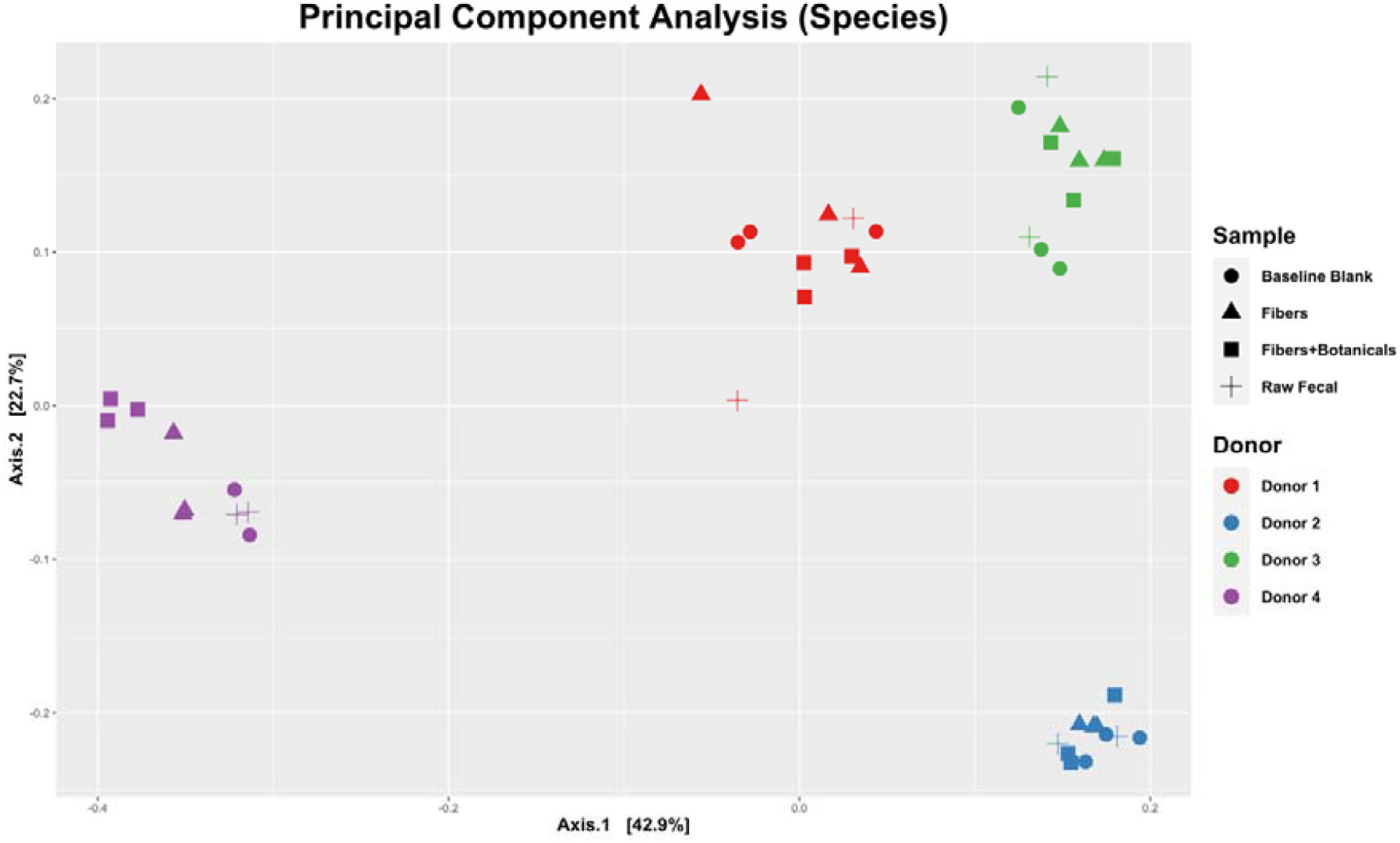
Principal components of analysis (PCoA) plot displaying UniFrac beta-diversity relationships among fecal samples, controls, and fermentations by donors.

**Figure 4:**
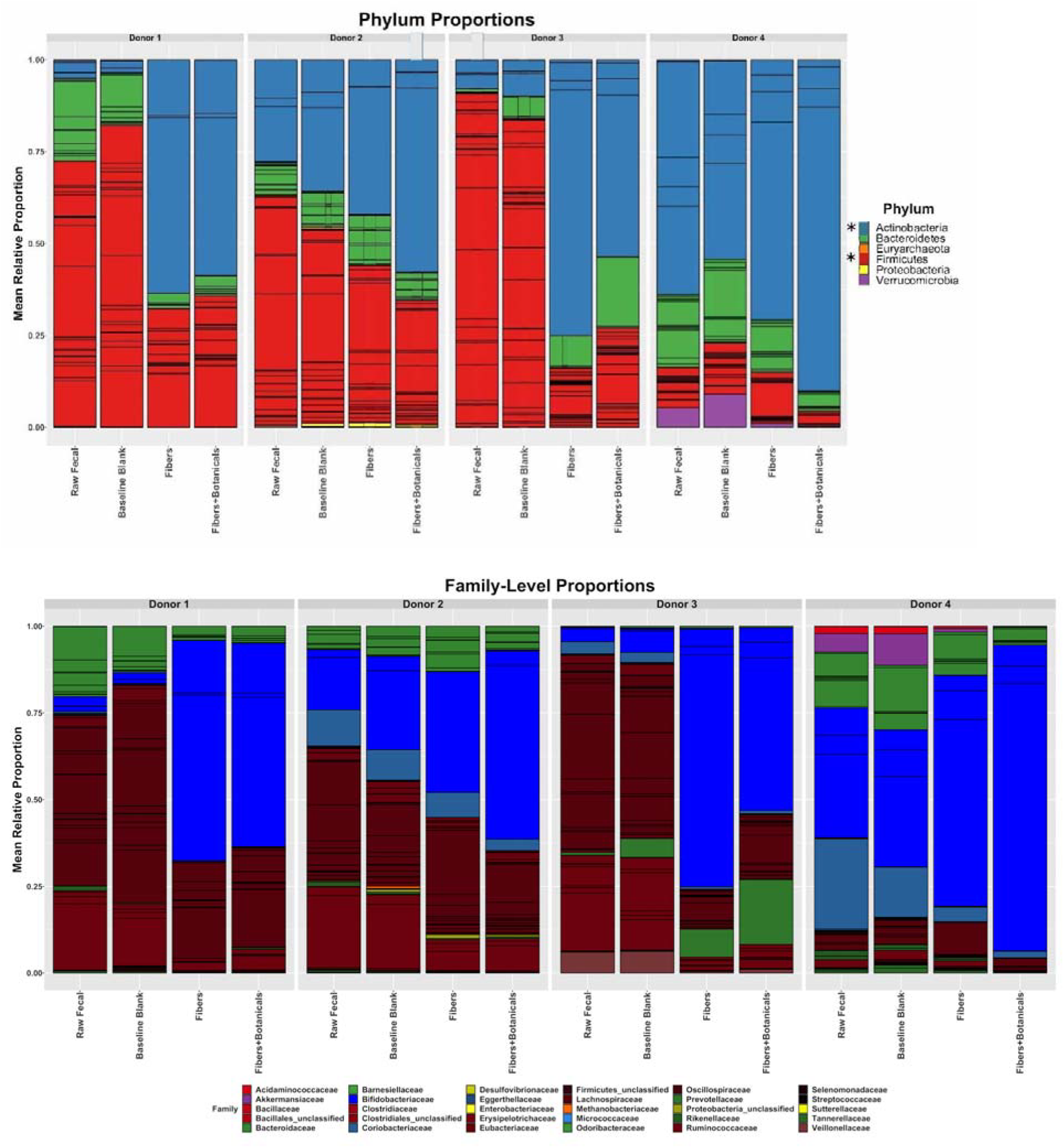
Relative abundances of metagenomic reads from fecal samples, baseline blanks, and fermentations, assigned by bacterial families. Individual OTUs are represented by the internal bars within a family’s color. An asterisk (*) beside legend symbols in panel A denotes significant donor condition effects (p < 0.001, ANOVA) in the same time-dependent direction for all 4 donors.

**Figure 5:**
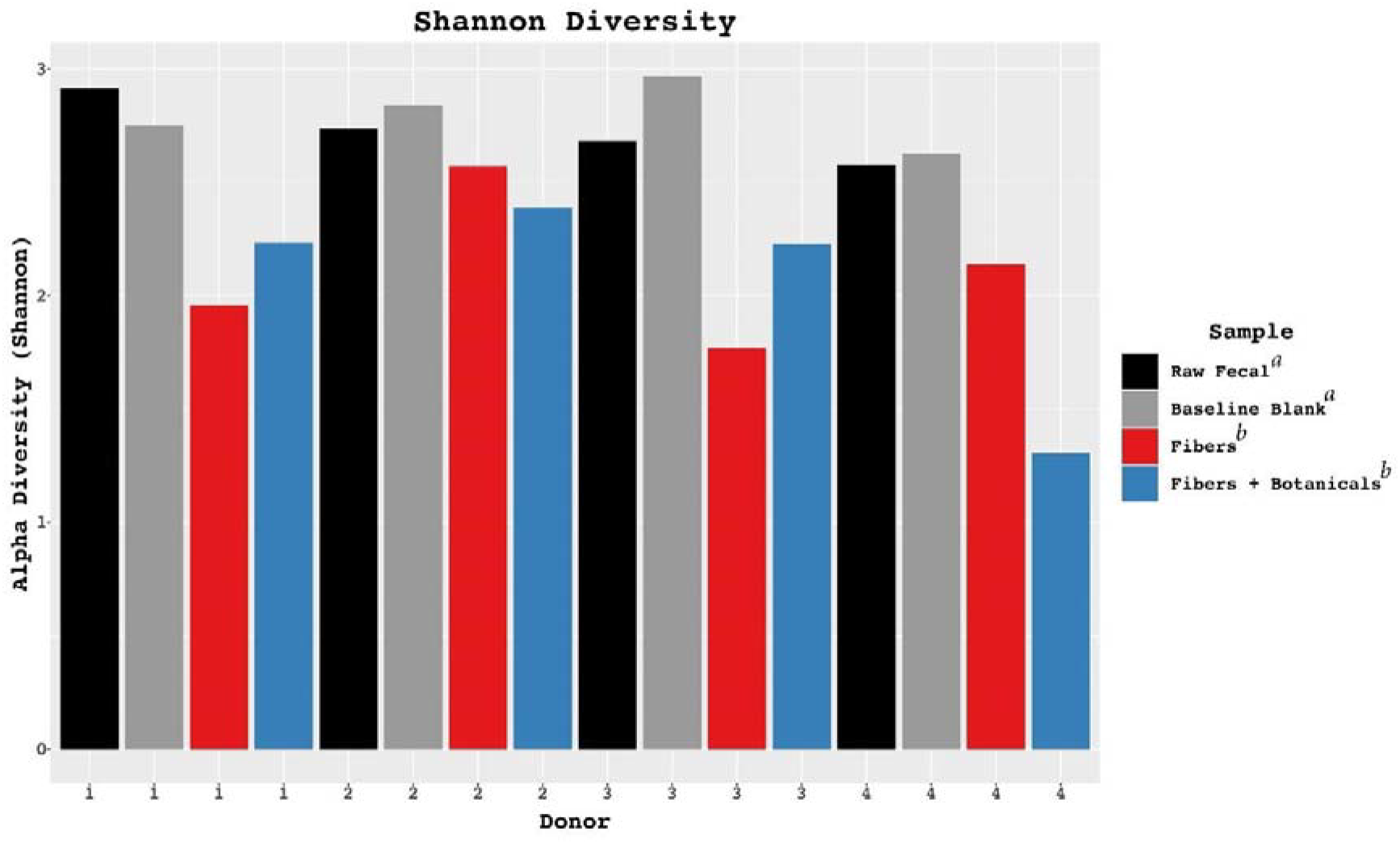
Alterations in Shannon diversity across donors in controls and fermentations compared with fecal samples. Letters *a* and *b* in the figure legend represent significantly different groups based on ANOVA results.

Mapping of bifidobacterial reads to the species level revealed strong, significant expansion of *B. adolescentis* populations, and this species was the most abundant bifidobacterial species after fermentation of OLISMART fibers, both with and without botanicals, in fermentations using all donors’ microbiota (Figure 6). Interestingly, the magnitude of the bifidogenic effect and the species selected were individual and dependent in some cases upon botanicals. With donor 1 and 3 microbiota, the initially small *B. longum* population was substantially increased after fermentation of OLISMART fibers, with and without botanicals. In contrast, this species was already more abundant in donor 2 and 4 microbiota pre-fermentation but was not increased (in relative abundance) by fermentation. Further, donor 2 and 3’s microbiota showed time-dependent increases in *B. pseudocatenalatum*, with donor 2’s *B. pseudocatenalatum* showing significant (p < 0.05, ANOVA) increases from timepoint 0 in only the botanical supplemented media while donor 3 saw significant increase in both conditions post-fermentation. *B. longum* showed a significant time-dependent increase (p < 0.001, ANOVA) in both donor 1 and 3, with donor 3 also exhibiting a significant increase in abundance in the presence of botanicals versus fiber-supplemented media alone (p < 0.001, pairwise t-test). Interestingly, in donor 2 fermentations the *B. longum* population was outcompeted by *B. adolescentis* when fermenting OLISMART fibers alone but was retained (and the entire bifidobacterial population increased) when botanicals were present. Donor 4’s initially large population of bifidobacteria contained sizable amounts of *B. adolescentis, B. longum*, and *B. bifidum*; in this case, only *B. adolescentis* markedly increased in relative abundance post-fermentation (p < 0.01, ANOVA) and the magnitude of this improvement was further increased in the presence of botanicals (p < 0.01, pairwise t-test). Less abundant bifidobacterial populations were influenced by botanicals in donor-specific ways. For example, in fermentations with donor 1’s microbiota, only fermentations containing the OLISMART fibers and botanicals resulted in the detection of *B. pseudocatenulatum*. Together, these data suggest that relative growth rates of bifidobacterial species are governed by donor context and the presence of botanicals in the fermentation.

**Figure 6:**
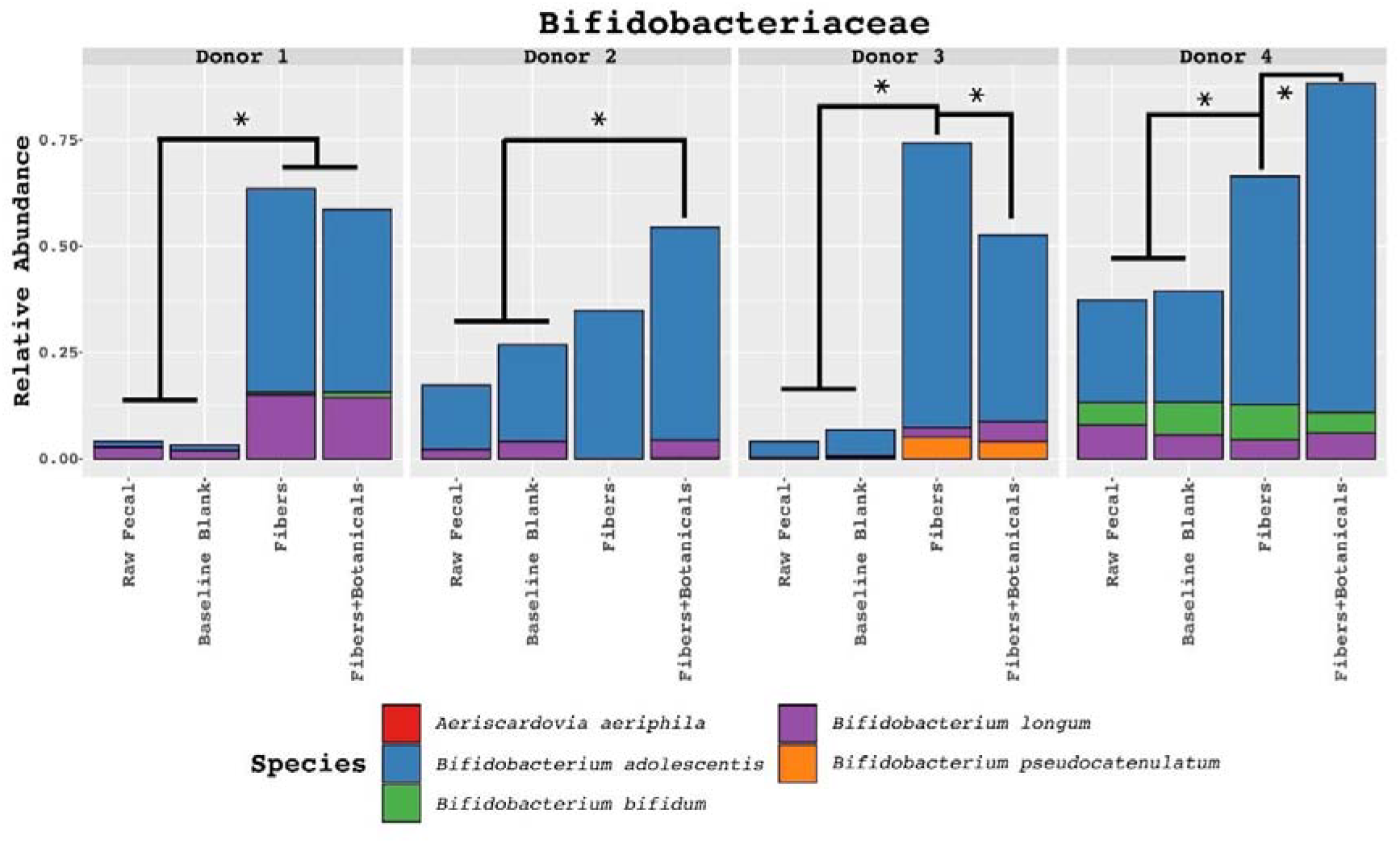
Relative abundances of metagenomic reads derived from fecal samples, controls, and fermentations assigned to family *Bifidobacteriaceae* and classified to the species level. Colored boxes in top-left of each facet represents significant fiber-dependent abundance increase on the species level. Statistically significant differences determined by ANOVA pairwise comparisons test (read-count as dependent and media type as the independent variable, split by donor) with p < 0.001. Symbol style: p-value < 0.001 (*).

## 4 Discussion

Fermentation of dietary components and their gut microbiome and metabolome effects are often investigated in isolation, whereas product formulations often contain blends of several functional ingredients. For example, research has been separately conducted on the beneficial effects of prebiotic fibers (Makki *et al*., 2018; Qin *et al*., 2023) and to a lesser extent for botanicals (Cefalu, Ye and Wang, 2008; Choi *et al*., 2020; Xu *et al*., 2022) but, to our knowledge, no research is published on the inter-related effects of both functional ingredient categories. To this end, we investigated this concept by employing OLIPOP, a widely consumed functional soda containing a mixture of 3 fibers and 4 botanicals, in *in vitro* fermentations of the prebiotic fibers in OLIPOP with and without the paired botanicals. As established previously (Yao, Chen and Lindemann, 2020) inter-replicate measurement differences in our proof of concept study were very small, suggesting the *in vitro* system employed here was technically robust for reproducible fermentation by gut microbiota.

Across all 4 donors employed, independent of initial community size and composition, a strong bifidogenic response and concomitant elevated SCFA production levels were established in fermentations containing the fiber mixture, either in the presence or absence of the botanical blend. Importantly, at the species level, the bifidobacterial community composition displayed subtle interpersonal differences in fermentation responses. This is a common observation and not surprising considering the fact that individuals harbor distinct endogenous bifidobacterial populations (Voreades, Kozil and Weir, 2014; Arboleya *et al*., 2016; Tierney *et al*., 2022). The consistency of the bifidogenic effect might be aided by the presence of multiple fibers in the blend, which may differentially stimulate the growth of distinct species. Hence, in addition to the general benefit of elevated SCFA levels triggered by fiber fermentation by any bifidobacterial species, species-specific health effects of bifidobacteria may play a complementary, donor-dependent role. To this end, fermentation of the fiber mixture by fecal microbiota resulted in increased levels of *B. longum, B. adolescentis*, or *B. pseudocatenulatum*, but not to the same extent in all donors. Recent studies suggest that *B. adolescentis* might be beneficial against constipation (Wang *et al*., 2017) and support kidney stone management (Abratt and Reid, 2010). Furthermore, it has been hypothesized that *Bifidobacterium* species have anti-obesogenic properties. While our study did not seek to measure the impact of adding live bacteria to assess obesity protective benefits, earlier work suggested potentially supportive effects when ingesting *Bifidobacterium pseudocatenulatum* strain CECT 7765 in a high-fat diet-fed obese mice model (Moya-Pérez, Neef and Sanz, 2015). They report addition of *B. pseudocatenulatum* resulted in lower serum cholesterol, triglyceride, and glucose levels and also reduced insulin resistance and positively impacted glucose tolerance. Given the significant global rise in obesity, prediabetes and type 2 diabetes rates, the observed increase of *B. pseudocatenulatum* in our study warrants further investigation to assess translational effects in human clinical studies using the botanical and prebiotic fiber blend from this study. Similarly, *B. longum* has been reported to protect against inflammatory bowel disease (Yao *et al*., 2021). Taken together, this suggests that while prebiotic fibers, such as those employed in this study, are likely to broadly elicit health benefits across individuals, these benefits may be personalized by individuals’ unique microbiome communities.

Recently, consumer food and beverage or supplements have evolved from containing single to multiple prebiotic and/or fibers components. However, research efforts to assess the interactive microbiome or health effect of multiple microbiome-active fibers is still in its infancy. Inulin is one of the most widely utilized prebiotic fiber substrates in functional foods and beverages, as well as supplements, principally due to favorable cost structure, substantial scientific research and compliant global regulatory position. Few *in vitro* or *in vivo* studies have examined the interactive beneficial microbiome or human health effects of combining inulin with other prebiotic or fiber substates, though these interactions are likely to be very influential on their behavior in the gut. Koecher et al., examined fermentation profiles both *in vitro* and in humans for fructo-oligosaccharides (FOS), inulin, gum acacia, and pea fiber alone or blended (Koecher *et al*., 2014). Additionally, Lecerf et al. sought to examine the differential effects of either xylo-oligosaccharide (XOS) alone or inulin-and-XOS mixture in a small human cohort (Lecerf *et al*., 2012). Interestingly, in this study XOS or XOS plus inulin did exhibit microbiome differences, yet crucial physiological effects were also observed when combinations were employed. Both of these studies indicate potentially beneficial effects of combining inulin and other prebiotic fibers ranging from increased SCFAs (Koecher *et al*., 2014) to attenuation of pro-inflammatory responses (Lecerf *et al*., 2012). Combinations of inulin and resistant dextrin, which was used in the present study, have not been previously investigated for microbiome or digestive health benefits. Cai et al. examined a milk powder with or without inulin and resistant dextrin blend on markers for type 2 diabetes (Cai *et al*., 2018). Studies combining inulin, resistant dextrin, and botanicals or, more broadly, any botanical and prebiotic fiber blends do not exist to our knowledge in the peer-reviewed scientific research. These prior studies and our findings indicate that products including prebiotic fiber blends need to be investigated to complement beneficial health impacts reported for isolated prebiotic fibers. Blending of fibers with other functional components such as botanicals could also lead to interaction effects, which need to be considered when investigating potentially beneficial microbiome and metabolomic impacts. Our proof-of-concept study suggests a potential need for the inclusion of study arms for each of the possible combinations of functional ingredients.

## 5 Conclusion

We sought to determine whether interaction effects of a functional blend would arise by conducting an *in vitro* batch culture fermentation experiment utilizing multiple donor’s fecal microbiota. Despite the fact our study encompassed gut microbial communities from a modest set of donors, it argues for the design of future clinical studies investigating the effects of blends of microbiome-active components to contain study arms investigating individual components and the full functional blends to measure potential interaction effects. Food-as-medicine interventions and dietary recommendations policy indicate that providing patients with medically tailored meals enabling achievement of daily recommended fiber intake led to beneficial health outcomes across multiple chronic conditions (Graff *et al*., 2023; Vedantam *et al*., 2023). As such, our study provides a roadmap for the functional beverage space and beyond, to rationally design studies and formulate products that result in beneficial health or microbiome impacts, with the ultimate aim to generate functional foods that bridge the fiber gap.

## 6 Acknowledgements

We cordially thank the Diversigen team for their help with the DNA isolation, sequencing and generation of the raw metagenomics data. We further wish to acknowledge the input of Renee Korczak, PhD, RD from OLIPOP for input and review of the manuscript. We also wish to acknowledge Sophie Pecher (Purdue University) for providing technical laboratory assistance and assay execution.

## 7 Conflict of Interest

PAB and NV consult for OLIPOP and receive compensation in turn. They did not accumulate the data nor perform the data analyses. SRL serves on the health advisory board of OLIPOP.

## 8 Author Contributions

NV and SRL conceived the study and directed the overall project. DGD carried out the experiments and, together with SRL, performed the data analyses, visualization and interpretation. DGD, PAB, NV, and SRL wrote the manuscript.

## 9 Funding

This study received funding and in-kind contribution of fibers and botanicals from OLIPOP, Inc. The funder had the following involvement with the study: design, writing and editing of the manuscript. BG is an employee, NV and PAB are paid consultants of OLIPOP, Inc. SRL serves on the Health Advisory Board of OLIPOP, Inc.

## 10 Data Availability Statement

The metagenomic datasets for this study can be found in the NCBI BioSamples Repository: using accession number PRJNA950354.

## Notes

https://www.ncbi.nlm.nih.gov/bioproject/PRJNA950354

### Summary of Updates

After rejection we have updated the format to align with a new journal we want to submit to

